# Harnessing biological variability for mechanistic inference: a stochastic framework applied to neural stem cell dynamics

**DOI:** 10.64898/2026.01.22.701043

**Authors:** Ren-Yi Wang, Diana-Patricia Danciu, Filip Z. Klawe, Anna Marciniak-Czochra

## Abstract

Inter-individual heterogeneity is often treated as noise, yet its temporal evolution can reveal regulatory mechanisms hidden from mean-field behavior. We present a stochastic framework that exploits variability for mechanistic inference in cell population dynamics. Using adult neurogenesis as a case study, we develop a state-dependent stochastic model of transitions between quiescent and active states and derive a diffusion approximation for the dynamics of both mean and variance. Applied to repeated cross-sectional data from wild-type and interferon-receptor knockout mice, we show that distinct regulatory mechanisms can produce similar mean dynamics but different fluctuation patterns. Jointly fitting mean and variance identifies proliferation-rate regulation as the dominant contributor to variability, while activation and self-renewal primarily govern average and long-term dynamics. Wild-type mice exhibit regulation of all three processes, whereas knockout mice lose activation control. These results show that population-level variability provides mechanistic information beyond average dynamics and helps distinguish between competing mechanistic models.

## Introduction

Mathematical models are widely used to infer regulatory mechanisms governing biological systems. A central challenge in this endeavour is non-identifiability: different regulatory hypotheses may reproduce the same experimental observations and therefore cannot be distinguished based on average population dynamics alone. This limitation often arises from the amount and granularity of available data. While numerous studies have successfully used mathematical models to uncover mechanisms underlying cell population dynamics, they have primarily focused on mean-field behaviour. To our knowledge, however, one layer of information that has not been systematically exploited so far is the temporal evolution of inter-individual variability arising from heterogeneous ageing. We propose that such variability may itself contain mechanistic information. Indeed, evidence for reproducible age-dependent trends in inter-individual variability has been reported across a range of biological systems ^[1,2,3,4,5]^.

To address this gap, we present a mathematical approach based on a diffusion-type approximation for analyzing the dynamics of population-level variability. Using adult neurogenesis as a case study, we demonstrate how this information can be exploited to infer regulatory mechanisms in neural stem cell populations. This approach enables the quantitative characterization of both the deterministic mean-field behavior ^[6]^ and higher-order variability, such as variances, of key system variables. Our theoretical framework is based on functional limit theorems ^[7]^, which provide a rigorous link between deterministic mean-field dynamics and stochastic fluctuations. The resulting diffusion approximation is conceptually equivalent to the Chemical Langevin Equation (CLE) and the Linear Noise Approximation (LNA), which are widely used in physics and systems biology^[8,9,10,11,12]^. The distinction lies not in the resulting diffusion limit, but in its derivation: whereas CLE and LNA are typically obtained through physical arguments and system-size expansions, our framework follows from functional limit theorems for density-dependent Markov processes^[7]^. More generally, the idea of exploiting stochasticity to alleviate non-identifiability has been discussed previously^[13]^. However, many existing stochastic inference approaches are designed for longitudinal observations and are therefore not readily applicable to the setting considered here. In particular, LNA-based filtering methods typically assume dependent longitudinal measurements from the same individual, whereas many experimental datasets consist of independent cross-sectional observations collected at predefined time points. Likewise, simulation-based stochastic approaches ^[14]^ are often computationally demanding and related existing parameter estimation routines are primarily designed to capture instantaneous fluctuations over relatively short time scales rather than the macroscopic evolution of variability ^[15]^. To overcome these limitations, we link the evolution of population distributions using limit theorems and the Bhattacharyya distance^[16,17]^. A key novelty of our approach is the sensitivity analysis of fluctuation magnitudes with respect to model parameters, allowing observed inter-individual variability to be directly related to model structure. This enables variability to be used to discriminate between competing regulatory hypotheses and identify underlying non-linear feedback mechanisms. In this way, stochasticity becomes not merely a source of noise but a source of mechanistic information that can be used to propose candidate regulatory mechanisms and infer underlying nonlinear dynamics.

To illustrate the applicability of our framework for mechanistic inference, we focus on adult neurogenesis as a case study. A hierarchy of mathematical models has recently been developed for this system, yielding several competing regulatory hypotheses that reproduce the available experimental data equally well. Adult neurogenesis therefore presents a clear identifiability challenge and provides an ideal setting to investigate whether population-level variability can improve model discrimination and reveal additional information about the underlying regulatory mechanisms. We complement this biologically motivated application with a toy model that highlights the general principle underlying the proposed framework.

Adult neurogenesis is the process through which mature neurons are generated from neural stem cells (NSCs). Adult NSCs can reside in either a quiescent (qNSC) or active (aNSC) state. Upon activation, qNSCs enter the cell cycle and may either self-renew or differentiate into neuronal progenitors. Experimental evidence has shown that neural stem cells and their progeny decline with age ^[18,19,20,21,22]^, contributing to age-associated cognitive impairment ^[23,24,25,26]^. Understanding the mechanisms regulating these population dynamics is therefore of considerable biological interest.

Previous mathematical models have progressively refined our understanding of adult neurogenesis. Early compartmental ODE models identified the transition from quiescence to activation as the primary regulatory layer governing neural stem cell maintenance ^[21,22]^. Subsequent studies comparing wild-type (WT) and interferon receptor knockout (IFNAGR KO) mice showed that perturbations of the activation process shift the regulatory balance towards self-renewal and differentiation, revealing a second layer of regulation ^[27]^. More recently, nonlinear feedback models revealed several alternative regulatory motifs that reproduce the available experimental observations equally well^[28]^.

These observations motivate our focus on the evolution of population-level variability. First, inter-individual heterogeneity decreases with age, such that older mice become more similar to one another than younger mice. Second, the nonlinear models for NSC dynamics^[28]^ have shown that multiple feedback hypotheses provide almost identical dynamics. In the following, we apply our framework to data from WT and IFNAGR KO mice comprising measurements of total NSC counts and active NSC fractions at different ages. Each data point corresponds to a different mouse, as they need to be sacrificed for the quantification. Consequently, the data represent repeated cross-sectional observations rather than individual trajectories. As shown in Fig. 1C, both datasets exhibit an age-related decline in total NSC counts, whereas the active fraction decreases in WT mice but increases in the IFNAGR KO setting.

**Figure 1:**
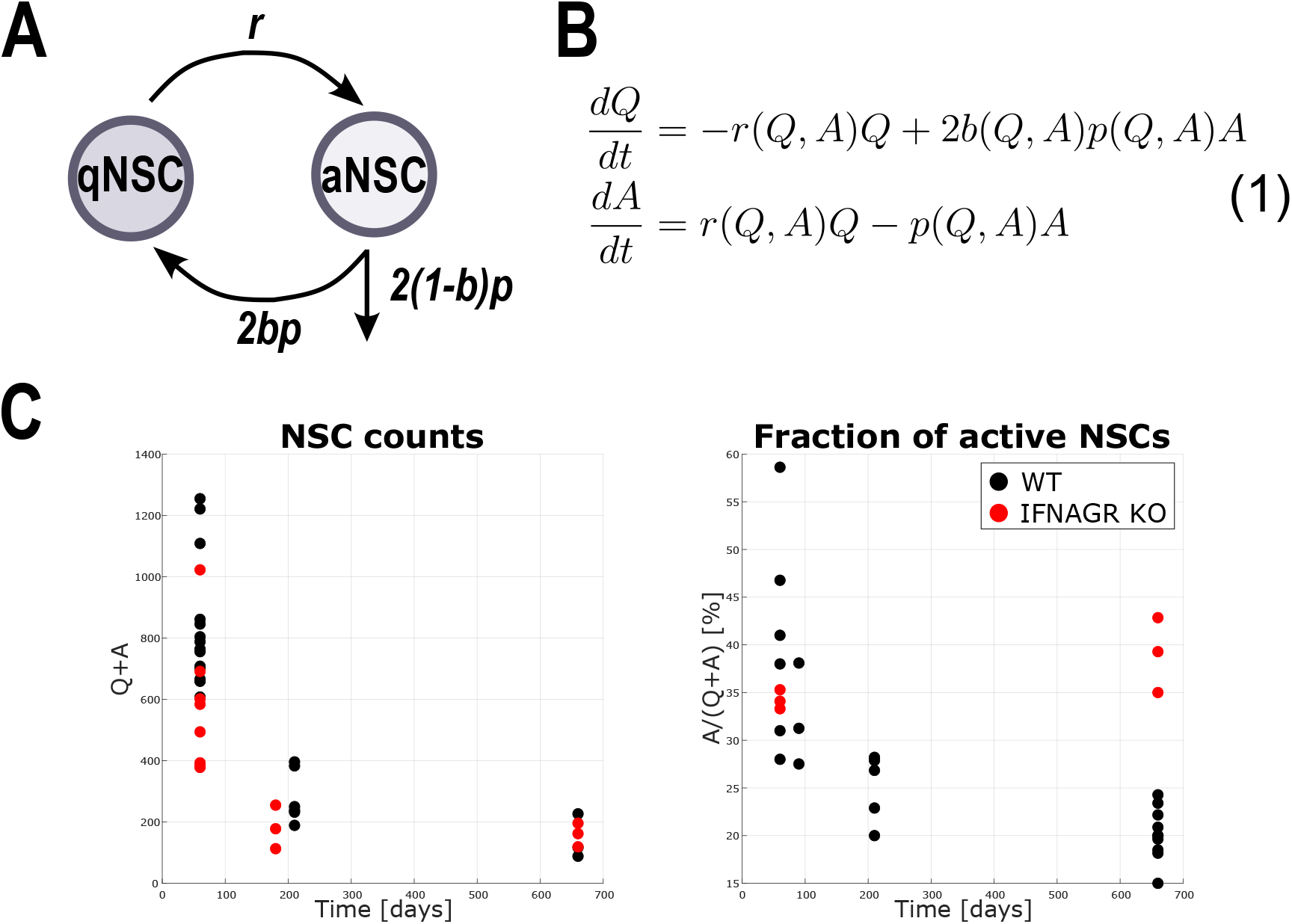
Conceptual overview of the study. **(A)** Schematic of compartmental model. A qNSC activates at a rate *r* and becomes an aNSC. An aNSC proliferates at a rate *p* and produces two qNSCs with a probability *b*, or exits the system by differentiating. **(B)** ODE system describing the dynamics depicted in panel **(A). (C)** Experimental data recording total number of NSCs (*Q* + *A*, left) and the fraction of aNSCs (*A/*(*Q* + *A*), right). The data comes from two experimental settings: wild-type (WT) mice, and genetically perturbed mice that had their Interferon *α* − *γ* receptors knocked-out (IFNAGRKO). This perturbation prevents the cells from sensing IFN inflammatory signals.

The remainder of this paper is structured as follows. We first present the stochastic framework and compare the resulting dynamics of population-level heterogeneity with experimental observations within the regulatory feedback class proposed by Danciu et al (2025) ^[28]^. We then illustrate the central principle of the framework using a toy model in which competing regulatory scenarios generate identical mean-field dynamics but distinct fluctuation patterns. Finally, we show how variability patterns can be exploited to discriminate between competing regulatory hypotheses and to elucidate the roles of different feedback mechanisms in shaping both mean-field behavior and inter-individual heterogeneity.

## Results

### Stochastic reaction network formulation captures lineage population dynamics

To investigate whether population-level variability contains additional mechanistic information, we formulate a stochastic model that describes the NSC populations qNSCs *Q*(*t*) and aNSCs *A*(*t*) as random variables with distributions determined by the following transitions (as depicted in Fig. 1A):

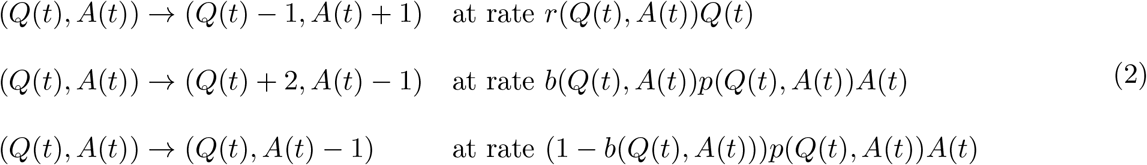

Here, a qNSC activates at a rate *r*(*t*) and an aNSC proliferates at a rate *p*(*t*). When an aNSC proliferates, it either produces two qNSCs with probability *b*(*t*) or differentiates with probability 1 − *b*(*t*). The time evolution of the parameters is considered as governed by non-linear feedbacks among cell populations. For simplicity, we will use *r*(*t*) := *r*(*Q*(*t*), *A*(*t*)), *b*(*t*) := *b*(*Q*(*t*), *A*(*t*)), *p*(*t*) := *p*(*Q*(*t*), *A*(*t*)). We apply a mathematical approach introduced by Wang et al (2025) ^[29]^ to analyze the asymptotic dynamics of population-level variability, grounded in a diffusion-type approximation for large initial populations, as found in the experimental data.

To facilitate reproducibility and adaptation of the proposed framework to related biological systems, we present the stochastic reaction network formulation and governing equations step by step.

Let *Q*^(*k*)^(*t*) and *A*^(*k*)^(*t*) denote the number of qNSCs and aNSCs at time *t* with initial population vector 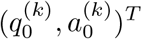, where *k* is a scaling parameter and elements in the initial population vector tend to ∞ as *k* → ∞. With appropriate scaling, we show by the functional law of large numbers (FLLN) in Lemma 1 that for large *k*, the scaled cell dynamics has the following mean-field approximation:

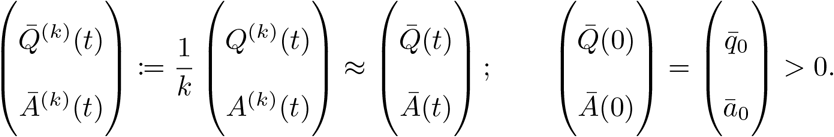

The deterministic FLLN limit 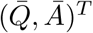 is characterized as the solution to an autonomous system of ODEs, representing the mean-field behavior of the dynamics, see Suppl. S2. This corresponds exactly to concentrations of cells described by the ODE system introduced by Kalamakis et al (2019) ^[21]^ and extended by Danciu et al (2025) ^[28]^. We may view *k* as a measure of physical space where the NSC dynamics take place. The mean-field scaled dynamics 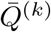 and 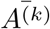 can thus be interpreted as the density of cell in the space. Hence, if signaling molecules are produced by NSCs, the mean-field scaled population is a measure of ligand concentration, a notion used in deriving the Hill equation in biochemistry^[30]^.

To additionally characterize stochastic deviations from the mean-field dynamics, we center and scale the concentrations around the mean by defining:

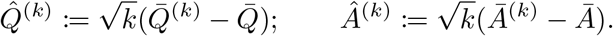

By the functional central limit theorem (FCLT) in Lemma 2 (Methods), the scaled deviations from the mean-field dynamics for large k,

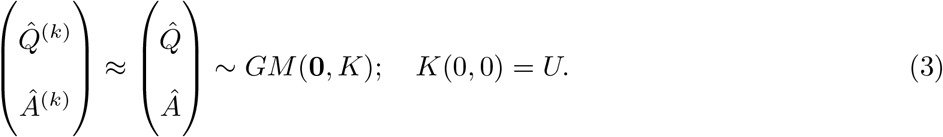

have a limit that follows a Gauss-Markov process^[31]^ with initial covariance matrix *U*. This process can be completely characterized by its mean function, which is identically zero in this case, and autocovariance function *K*(·,·).

Consequently, the dynamics of the concentrations of quiescent and active NSCs is given by the superposition of a deterministic central tendency and appropriately scaled stochastic fluctuations

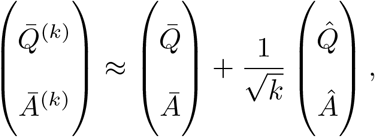

with dynamics governed by ordinary and stochastic differential equations, respectively:

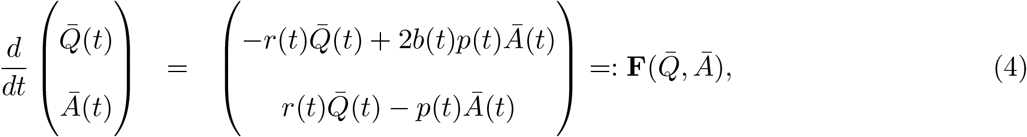

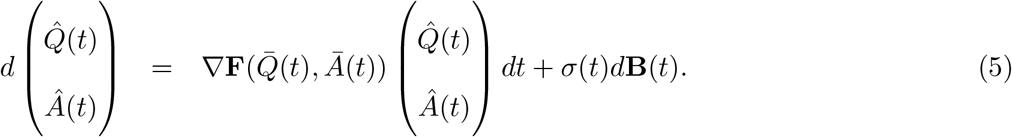

The time-dependent coefficient *σ*(*t*) arises from the application of the Poisson FCLT^[7,32,33]^ (see Suppl. S3 for a heuristic explanation). Analogous to the classical central limit theorem, we need to derive the limiting distribution, characterized by its mean and variance, for the centered and scaled random variables. In the stochastic process limit, we center and scale the Poisson processes to obtain the diffusion limit, which is characterized by its drift parameter 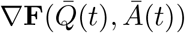 and diffusion coefficient *σ*(*t*). The diffusion coefficient is given by the rates of the actions that the variables *Q* and *A* can perform and the respective changes (a detailed derivation is found in Methods).

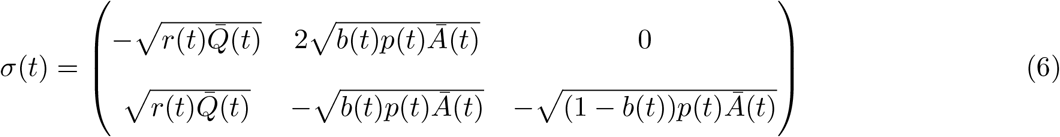

The variance function for 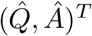 is characterized by

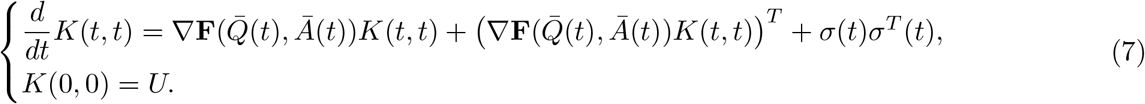

### Model inference reveals insights into biological mechanisms

We previously derived equations governing the mean-field dynamics (4) and the behavior of stochastic deviations from the mean-field (5). In the next step, we estimate parameters in order to use the information from the fluctuation dynamics to gain new insights into the underlying mechanisms of the neural populations. As our data record total NSC counts and fractions of aNSCs, a first step is to perform a coordinate transformation:

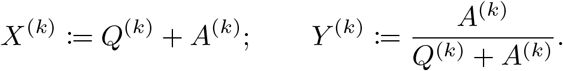

and derive their corresponding FLLN and FCLT limits. By continuous mapping and the delta method (Theorem 1 and 2, Methods), we show that for large *k*,

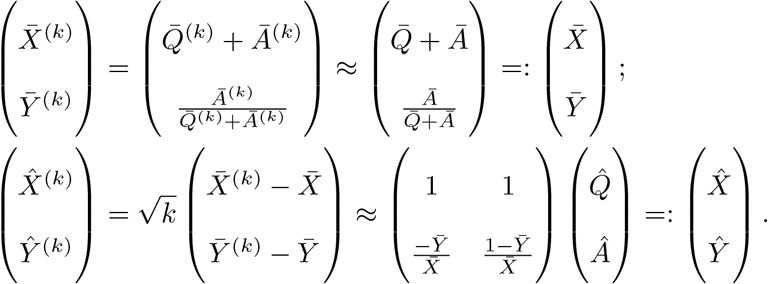

This transformation is required both to estimate the parameters of the model and to express the resulting dynamics in the variables corresponding to experimentally observable quantities. Importantly, the proportion of aNSCs relative to the total population provides more informative insight into the system than the absolute cell count. To apply theses asymptotic results, we take *k* = 1000, which reflects the typical scale of the NSC population at 60 days of age.

In the following, we investigate the evolution of the stochastic system in a few scenarios comprising combinations of feedback mechanisms, extended from those proposed by Danciu et al (2025) ^[28]^. The activation rate *r*(*t*) is an increasing Hill-type function w.r.t. the density of cell types 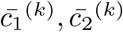 and the self-renewal probability *b*(*t*) is a decreasing Hill-type function w.r.t. 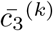 (8). The cells *c*_*i*_ correspond to linear combinations of variables *Q* and *A*. Additionally, we allow the regulation of the proliferation rate *p*(*t*) by the total concentration of NSCs, 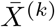, *via* a similar increasing Hill-type function. This choice is biologically motivated by the existence of paracrine signaling within the neural stem cell niche, where diffusible factors exchanged between NSCs and surrounding niche cells give rise to effective density-dependent feedback on proliferation dynamics^[34,35]^.

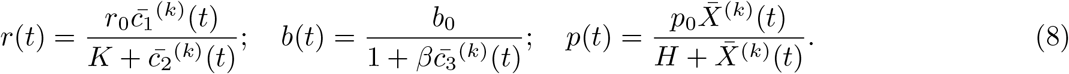

We explore, on the one hand, which feedback mechanisms are most likely out of the considered scenarios and, on the other hand, the regulation of which system parameters is essential.

Because the model describes stochastic population dynamics, fitting only mean trajectories or least-squares residuals would neglect information contained in the full cell-state distributions. We therefore formulate parameter inference as a comparison between experimentally observed and simulated probability distributions.

To quantify distribution similarity, we employ the Bhattacharyya distance^[36,16,17]^, which measures the overlap between probability distributions and is symmetric and numerically stable in the presence of stochastic variability. For (multi-)normal distributions it has a closed form depending solely on the mean and standard deviation of the distributions under comparison. Alternative divergences such as the Kullback–Leibler divergence or Wasserstein distance could also be considered; however, the Bhattacharyya distance provides a particularly natural measure for comparing noisy population distributions generated by stochastic chemical reaction network simulations.

Because the experimental data exhibit slight skewness while the stochastic framework assumes Gaussianity, the data were first approximated by a log-normal distribution, and a Gaussian was then fitted to the high-density (main body) region obtained by symmetric truncation of the tails about the mode (see Methods and Suppl. Fig. S1). The model parameters were inferred by minimizing the Bhattacharyya distance between the data-derived and model-predicted Gaussian distributions, which as previously mentioned is fully determined by their respective means and standard deviations (see Methods). In the figures, red dots indicate data points that fall within the estimated Gaussian region, whereas black dots denote points that were excluded by the approximation procedure.

First, we observe that the various regulatory scenarios are equally capable of capturing the data dynamics well, when the same number of system parameters are regulated (Fig. 2A,B), similarly to results reported by Danciu et al (2025) ^[28]^. This supports the choice of regulatory functions (8), showing that not only is the mean-field behavior captured, but also the variability, regardless of which cell populations are involved in the regulations. Furthermore, from mathematical analysis we observe that in order to capture the decrease of cell concentrations, the self-renewal parameter *b <* 1*/*2. In this case, if self-renewal is not regulated (*b* constant), the system asymptotically converges to zero. Otherwise, with the Hill-type regulation, *b*_0_ = 1*/*2 is a bifurcation point such that either only the trivial steady state exists (*b*_0_ *<* 1*/*2) and it is stable or there are two steady states (*b*_0_ *>* 1*/*2): an unstable trivial and a stable positive one. As reported before^[28]^, it appears that if qNSCs are involved in inhibiting self-renewal (blue, red and purple lines in Fig. 2B) cell concentrations converge to a positive steady state, whereas scenarios in which only aNSCs inhibit self-renewal lead to extinction (pink and green lines).

**Figure 2:**
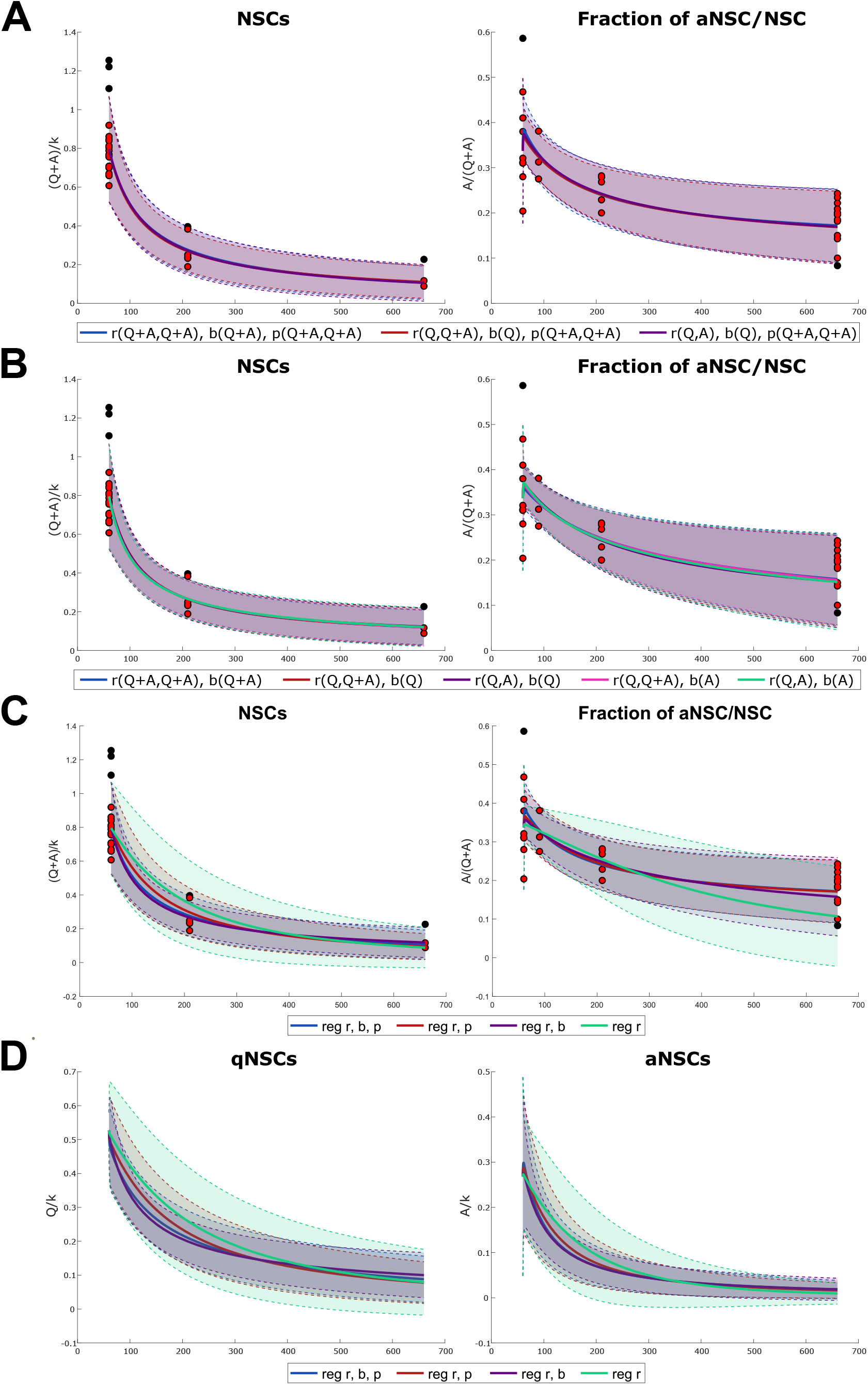
Comparisons among different regulatory feedback scenarios. We present results from models incorporating various types of feedback and different concentration variables appearing in the nonlinear terms (see Eq. (8), *k* = 1000). Solid lines denote mean behavior, while shaded regions indicate prediction intervals. The number of NSCs and the fraction of aNSCs are shown for: **(A)** regulation of activation, self-renewal probability, and proliferation via different cell concentrations; **(B)** regulation of activation and self-renewal probability via different cell concentrations; **(C)** comparison of models with different combinations of regulated parameters, through feedbacks governed by the entire NSC population, i.e. *c*_1_ = *c*_2_ = *c*_3_ = *X* in Eq. (8). Panel **(D)** presents concentrations of qNSCs and aNSCs for the same feedback configurations as in **(C)**.

Secondly, a new insight that we gain only by additionally accounting for the stochasticity is related to how the various system parameters influence the inter-individual heterogeneity (Fig. 2C). From previous work ^[21,22,28,37]^ we know that in order to explain the decrease of the fractions of aNSCs in time, the activation rate *r* has to decrease with age, and equivalently increase with cell numbers as in Eq. (8). Therefore we selectively allow certain parameters to be regulated while others are kept constant: cases [*r, b, p*] (blue), [*r, p*] (red), [*r, b*] (purple) and [*r*] (green) regulated. We observe that even though in all cases the mean-field behavior is captured well (when regulating *r*) or very well (when regulating at least one additional parameter), there are differences in how well the variability is fitted. We can see that only allowing the regulation of [*r*] and [*r, b*] leads to prediction intervals that are wider than the variability observed in the data, especially in the fraction of aNSCs. On the other hand, the cases of regulating [*r, p*] and all [*r, b, p*] lead to almost perfectly overlapping means and prediction bands, which very nicely recapitulate the data. Upon inspecting all these combinations of regulated parameters, our model suggests that the proliferation rate *p* is the main parameter that influences the amplitude of fluctuations and its regulation is crucial to capture the stochasticity in the data. Additionally, Fig. 2D shows similar plots for the original variables *Q* and *A* and suggests that the variability in qNSCs is much larger than that in aNSCs. The model thus indicates that while regulation of activation and self-renewal primarily shapes mean-field trajectories and long-term population maintenance, regulation of proliferation has the strongest effect on attenuating fluctuation amplitudes for the active fraction dynamics.

In a second data set, where the IFNAGR have been genetically knocked out, we observe an earlier decrease in NSC numbers and a slight increase in the fraction of aNSCs. Previous works ^[27,28]^ suggested that in this perturbed system the activation rate *r* does not decrease with aging, and that the fraction of self-renewal *b* gains a greater impact, counteracting the effects of an almost constant activation rate *r* on the depletion of NSCs. As such, in the scenario where the regulations are governed by the total population of NSCs (*c*_1_ = *c*_2_ = *c*_3_ = *X* in Eq. (8)), we look at a few cases of parameter combinations: [*r, b, p*] (blue), [*b, p*] (red) and [*r* (with *r*_0_), *b*] (purple). The latter case is a modified version of the feedback function for the activation rate 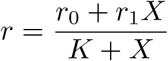, which was introduced by Danciu et al (2025) ^[28]^, and which allows capturing both the decrease in WT data and the increase in the IFNAGRKO data for the fraction of aNSC, whilst keeping the proliferation rate *p* constant. As before, all three cases considered are able to recapitulate the mean behavior, as shown in Fig. 3A. However, differences in the fit to the variability in the data can be observed, and the model suggests that having the activation rate *r* regulated is not essential to explain the data (Fig. 3B).

**Figure 3:**
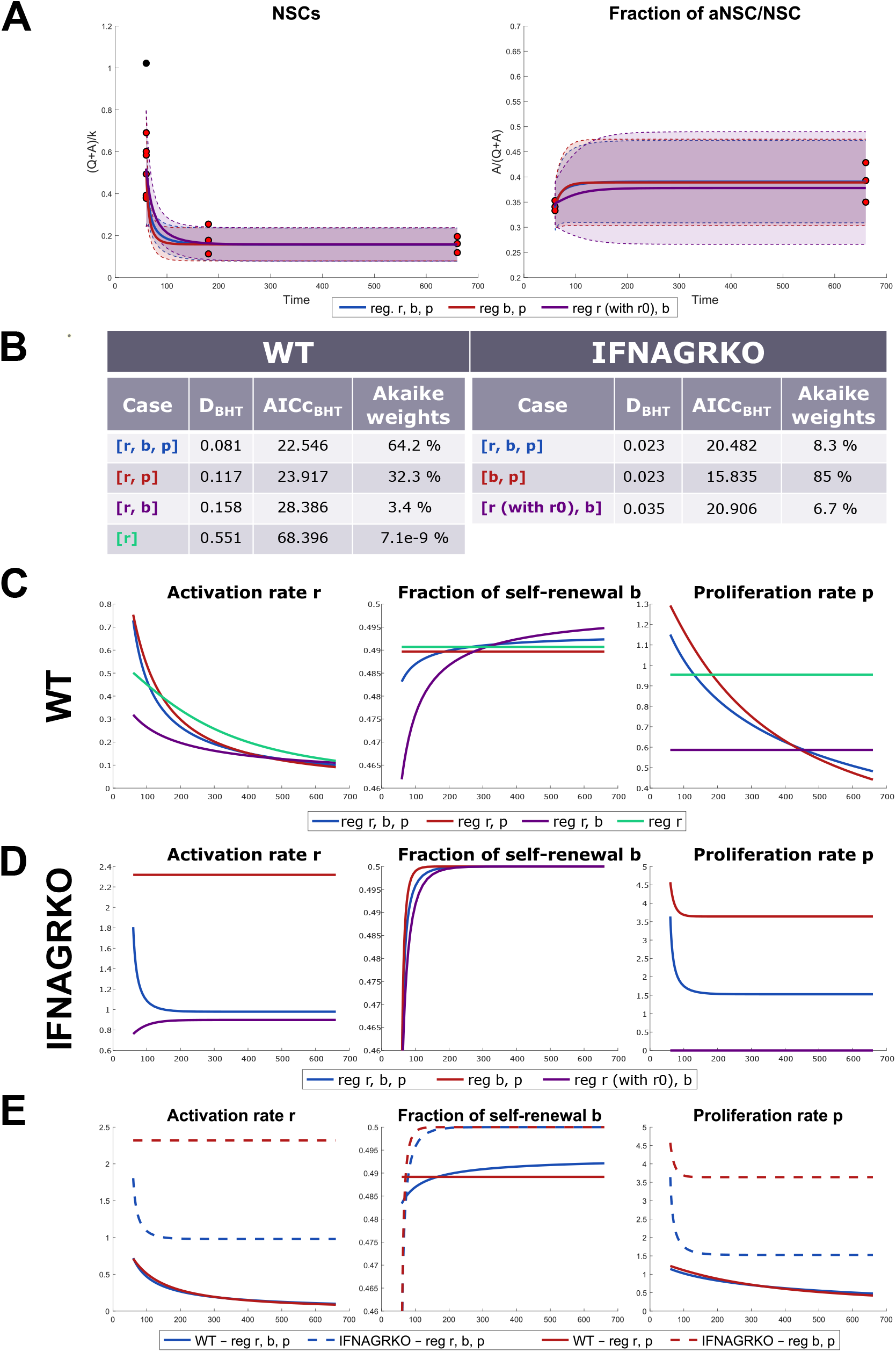
IFNAGRKO and parameter evolution. **(A)** Concentration of NSCs and fraction of aNSCs in IFNAGRKO under different combinations of parameter regulations, with *k* = 1000. Solid lines represent mean values, and shaded regions indicate prediction intervals. All feedback regulations are governed by the total NSC population. **(B)** AICc values and Akaike weights for all tested scenarios for WT and IFNAGRKO. Comparison of parameter evolution (activation rate, self-renewal probability, and proliferation rate) across different feedback mechanisms: **(C)** in WT and **(D)** in IFNAGRKO. **(E)** Direct comparison of parameter evolution between WT (solid lines) and IFNAGRKO (dotted lines), in the two best scenarios for each setting: blue - regulation of all three parameters [*r, b, p*], and red - regulation of proliferation and one extra parameter ([*r, p*] for WT and [*b, p*] for IFNAGRKO).

A next step is to inspect the dynamics of the system parameters *r, b, p*. In Fig. 3C–E, we show the time evolution of the parameters in the cases corresponding to Fig. 2C in WT, Fig. 3A in IFNAGRKO mice, and in a comparison between the two best cases for each experimental setting. In WT, we observe that in the two best cases (blue and red) the dynamics of the parameters is very similar. We thus learn that even tiny changes in self-renewal can have a big impact on the system. In the IFNAGRKO, a new insight is that regulating the proliferation rate *p* allows capturing the increase in the fraction of aNSCs even with a constant or decreasing activation rate *r* (Fig. 3). Finally, inspecting the parameter dynamics for the best two cases in the WT (solid curves) and IFNAGRKO (dotted curves) data sets, we find much larger values for all parameters in the perturbed setting (Fig. 3C-E).

Finally, by using a Bhattacharyya distance-informed score similar to the corrected Akaike Information Criterion (AICc) for model selection^[38,39,40,41]^ and its corresponding weights (see Methods), we gain insight into the parsimony of the various cases considered: AICc suggests the best-scoring model, and the Akaike weight indicates the likelihood of it being truly the best. The Akaike-like scores and weights for the WT data set support the case in which all parameters are regulated, with a likelihood of 64% (Fig. 3B, left), closely followed by the scenario with regulated [*r, p*] with a likelihood of 32 %. In IFNAGRKO data, the Akaike-like scores and weights select the case of regulating only [*b, p*] with a likelihood of 85% (Fig. 3B, right). Thus, although all three parameters [*r, b, p*] appear regulated in WT mice, the IFNAGRKO data strongly support a loss of regulation of the activation parameter *r* (Fig. 3B). This finding is consistent with experimental evidence implicating interferon signaling in the control of NSC activation and exit from quiescence, suggesting that removal of interferon receptors disrupts an important regulatory layer governing qNSC activation ^[27,42,43]^. Furthermore, results from both experimental settings support the regulation of the proliferation rate *p*, whereas there exists a tradeoff between regulating activation *r* and self-renewal *b* parameters.

### Stochastic framework can be adapted to diverse biological systems and data sets

While neural stem cell dynamics provide a biologically motivated application, the key idea underlying the framework is more general. To demonstrate this, we consider a minimal toy model in which competing regulatory mechanisms produce identical mean-field dynamics but distinct fluctuation patterns. This example isolates the role of variability and illustrates how the proposed approach can be applied beyond the specific context of adult neurogenesis.

Let us consider two Markov models:

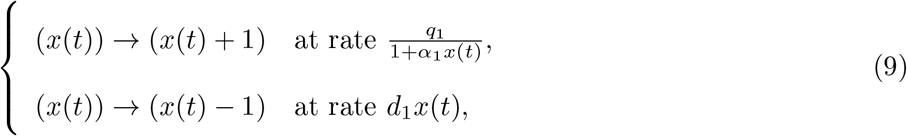

and

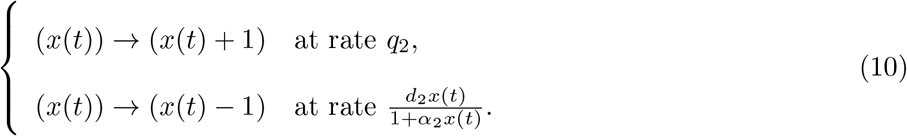

Both systems describe a population *x* (of individuals/cell types/chemical molecules/particles etc.) governed by the same mechanistic processes: inflow/production ∅ → *x*, and outflow/death *x* → ∅. The difference between models lies in the regulation of system parameters: the first model (9) has the production rate regulated through an inhibitory Hill-type function, while in the second model (10) the degradation rate is modulated by a similar feedback. The parameters *q*_1_, *q*_2_, *d*_1_ and *d*_2_ are positive constants, whereas *α*_1_ and *α*_2_ are non-negative.

As the scale of the system grows, we switch from tracking discrete particle numbers to a continuous concentration. In this limit, by the functional law of large numbers, the stochastic Markov process is well approximated by the following deterministic ordinary differential equations:

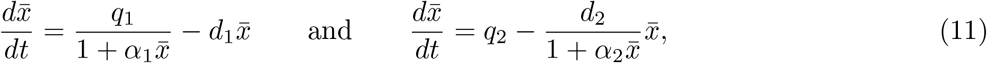

which govern the average time evolution of the concentration.

Let us take the following relation between parameters *α* := *α*_1_ = *α*_2_, *q*_1_ = *q*_2_(1 + *αx*_*s*_) and *d*_2_ = *d*_1_(1 + *αx*_*s*_), where 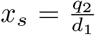. Then, it is clear that *x*_*s*_ is the only positive steady state to both ODEs from Eq. (11). Not only do both models, with the two different regulations, lead to the same steady state, but their mean-field dynamics is also very similar. However, an important difference between the two models is the pattern of fluctuations that they exhibit, particularly when ^1^ is much smaller than *x*_*s*_. Fig. 4 shows data generated from randomly choosing ten data points at each of seven time points out of hundreds of stochastic trajectories of the Markov models with regulated production (left column) and regulated degradation (right column). This way of generating the data captures the disadvantages of experimental data where animals have to be sacrificed and thus time-series are actually formed by numbers from different genetically equivalent individuals.

**Figure 4:**
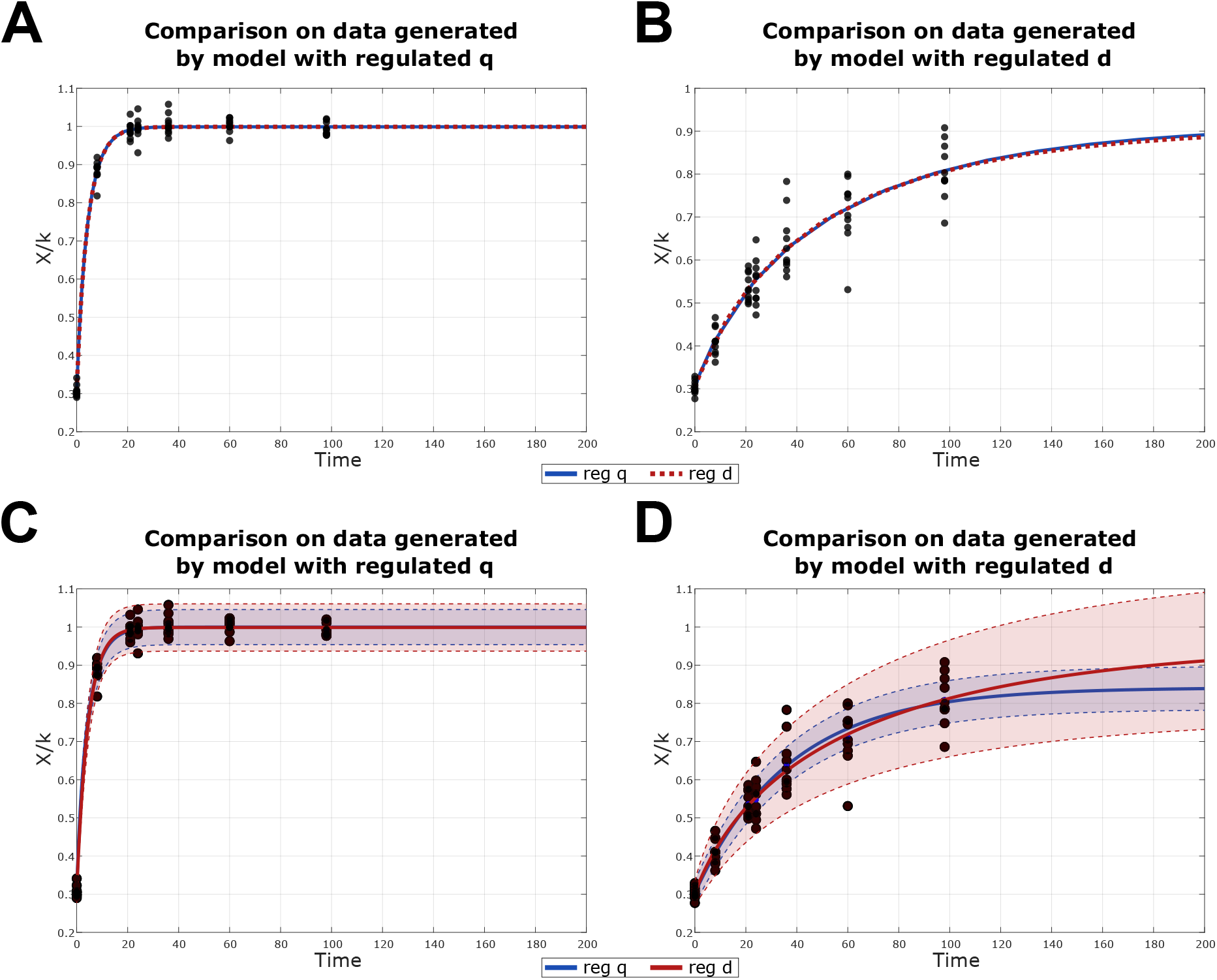
Toy model results. Fitting the models to data generated by Gillespie’s algorithm for *q*_2_ = 100, *d*_1_ = 0.1 and *α* = 0.01, by the two models: (9) with regulated *q* (left column) and (10) with regulated *d* (right column). To turn from discrete particle numbers to concentrations we rescale the system by k=1000. **(A)** ODEs fitted to the dataset generated by the model with regulated *q* (9). Akaike weights [49.902%, 50.098%] for models and (10), respectively. **(B)** ODEs fitted to the dataset generated by the model with regulated *d* (10). Akaike weights [49.493%, 50.507%] for models (9) and (10), respectively. **(C)** Result of fitting both mean and variance to the dataset generated by the model with regulated *q* (9). Akaike weights [99.996%, 3.98*e* − 03%] for models (9) and (10), respectively. **(D)** Result of fitting both mean and variance to the dataset generated by the model with regulated *d* (10). Akaike weights [4.21*e* − 25%, 100%] for models (9) and (10), respectively.

Taking the standard weighted least-squares approach of simply fitting the ODEs (11) for mean-field dynamics to the data generated by each Markov process (9) and (10), we are able to perfectly capture the average dynamics with both models. This is further emphasized upon computing their Akaike-like weights: both models are equally plausible, with respective weights of approximately 50%. Thus, despite the fundamentally different regulatory mechanisms, the two models are statistically indistinguishable when only mean-field dynamics are considered.

However, using the approach presented in this paper, we track not only the mean value but also the variability of the data. Fitting the parameters using equations corresponding to (4) and (7) with the Bhattacharyya distance, we are taking into account also the inter-individual heterogeneity. Note that in this toy model, the amplitude of fluctuations is an intrinsic property of the regulatory mechanism considered: even though the asymptotic behavior and initial conditions are the same, the speed of convergence and, more importantly, the magnitude of variability differ. Even though both models are equally able to capture the mean-field dynamics of both data sets, with no possibility of model selection (Fig. 4A–B), leveraging the additional information encoded in population-level variability allows us to correctly identify the underlying regulatory mechanism with more than 99% confidence (Fig. 4C–D).

This example shows that population-level variability can resolve ambiguities that remain hidden at the level of average dynamics and illustrates the broader applicability of the proposed framework. The main assumption of the method is the availability of sufficiently large populations and approximately Gaussian observations at each time point. Below we briefly summarize the workflow.

1. Given a biological system, start from a Markov model of the different populations with transitions among them based on biological knowledge and assumptions (see Fig. 1A).
2. Build stochastic model based on the assumed transitions (see Eq.(2))
3. Upon centering and scaling the concentrations around the mean, obtain:
  i. ODE for mean-field dynamics (4),
  ii. SDE for dynamics of fluctuations (5), with respective
  iii. ODE for variance dynamics (7), depending on the Jacobian of the mean field dynamics.
  iv. Approximate experimental data by Gaussian distributions.
4. Estimate parameters by comparing the data and model distributions using the Bhattacharyya distance for Gaussians (Methods), with closed form depending only on the mean and variance values.
5. Compute mean and 95% prediction intervals with the fitted parameters.

## Discussion

In the current paper, we have presented a practical stochastic framework that can be used for inferring additional mechanistic insights from heterogeneities in the experimental data. Often, the scarcity or resolution of data leads to non-identifiability issues, such that multiple competing hypotheses appear valid. Our stochastic method can improve this aspect by extracting an extra layer of information from the variability in the data.

While our results confirm the validity of the previously proposed class of Hill-type regulatory feedback mechanisms in the neural lineage ^[28]^, they also show that in this case study of adult neurogenesis, even when experimental fluctuations are taken into account, the specific cell types involved in the regulations cannot be uniquely identified. However, incorporating stochastic fluctuations enables discrimination between competing regulatory hypotheses at a different level. Specifically, we find strong support for models that additionally include regulation of the proliferation rate, which are indistinguishable from models with constant proliferation rate at the level of mean behavior, but are clearly favored when the temporal evolution of variability in the experimental data is taken into account. Existing biological studies support the regulation of proliferation through paracrine and autocrine signals, via pathways such as EGF, Notch or Wnt^[35,44]^. Our models also suggest that experimental perturbations targeting cell-cycle progression should produce larger changes in the magnitude of inter-individual variability than perturbations affecting activation alone. Additionally, we showed that while the regulation of the activation rate *r* and the fraction of self-renewal *b* parameters primarily influence the mean-field dynamics and the long-term steady states, the main regulation that governs the amplitude of stochastic fluctuations in the system is that of the proliferation rate *p*, in agreement with the conclusions from the model of hematopoiesis considered by Wang et al (2025) ^[29]^. Furthermore, we showed that regulating *p* in addition to self-renewal *b* is sufficient to capture the slight increase observed in the IFNAGRKO fraction of aNSC data set, and that the activation rate can remain constant. The models also suggest higher activation and proliferation rates in the perturbed system, as well as more pro-neurogenic differentiation (Fig. 3E), which was also broadly suggested by biological studies^[45]^. Model selection based on Akaike-like information criteria further suggests that this scenario is the most parsimonious explanation of the data, indicating that interferon receptor knockout suppresses activation regulation that is otherwise essential in the wild-type setting^[27]^. Our results demonstrate that variability dynamics can substantially reduce non-identifiability and improve discrimination between competing regulatory mechanisms. Nevertheless, in the neural stem cell system studied here, multiple feedback scenarios involving different regulatory cell populations remain consistent with the available data.

The presented stochastic approach is a practical, scalable, and versatile method that turns variability in the data into a source of information. As further illustrated on our proposed toy model, it can be adapted to a wide range of Markov-governed systems and experimental settings, including single-cell lineage tracing in stem cell compartments, clonal dynamics during tissue homeostasis and development, heterogeneous signaling and gene expression networks, or variable treatment responses in cancer cell populations. One major advantage of the approach is circumventing the need for Gillepsie-type stochastic simulations, which can be computationally expensive and for which the estimation of parameters that can capture the macroscopic evolution of fluctuations is not trivial. The close correspondence between the prediction intervals and multiple realizations of the stochastic model is shown in Suppl. Fig. S1. With our diffusion-propagation stochastic approach, parameter estimation consists solely of defining an appropriate objective function that uses a distance for comparing distributions, which can be minimized by available optimization algorithms, being thus straightforward and efficient. Incorporating information from the variability in the data as additional constraints for the optimization can further help reduce issues related to non-identifiability and thus improve model discrimination. More broadly, our results suggest that population-level variability should not be viewed merely as noise, but as an additional source of information for mechanistic inference whenever competing biological hypotheses produce similar mean-field behavior.

### Limitations of the study

- The ability of the framework to discriminate between competing mechanistic models depends on the quantity and informativeness of the available data. Although incorporating variability improves model discrimination, it may not uniquely identify the underlying mechanism in all settings.
- The current implementation assumes approximately Gaussian experimental distributions. For positive-valued measurements, suitable transformations or Gaussian approximations (e.g. of truncated lognormal distributions) can extend the applicability of the framework.
- The framework exploits information contained in the first two moments of the population dynamics. Incorporating higher-order moments or full distributional information may further improve discrimination between competing mechanistic models.

## Resource availability

### Lead contact

Requests for further information and resources should be directed to and will be fulfilled by the lead contact, Anna Marciniak-Czochra.

### Materials availability

This study did not generate new unique materials.

### Data and code availability

- This paper analyzes existing, publicly available data, accessible at https://doi.org/10.1016/j.cell.2019.01.040 and https://doi.org/10.15252/emmm.202216434.
- All original code has been deposited at Zenodo (https://doi.org/10.5281/zenodo.18399730) and will be publicly available as of the date of publication.
- Any additional information required to reanalyze the data reported in this paper is available from the lead contact upon request.

## Acknowledgments

The project was supported by the European Research Council (ERC) under the European Union’s Horizon 2020 research and innovation programme (synergy project PEPS, no. 101071786), and by the Deutsche Forschungsgemeinschaft (DFG) within the Collaborative Research Centre SFB1324 (B05 of AMC). R-YW acknowledges the support from the Herbert and Florence Irving Institute for Cancer Dynamics at Columbia University. The authors thank Prof. Marek Kimmel and Dr. Alexey Kazarnikov for their feedback and invaluable advice on manuscript preparation.

## Author contributions

R-YW: Method development, Mathematical analysis, Interpretation of the modeling results, Manuscript preparation - original draft and revisions;

D-PD: Model development, Algorithm implementation, Interpretation of the modeling results, Manuscript preparation - original draft and revisions;

FZK: Mathematical analysis, Model development, Interpretation of the modeling results, Manuscript preparation - original draft and revisions;

AM-C: Conceptualization, Supervision, Model development, Interpretation of the modeling results, Manuscript preparation - review and editing;

## Declaration of generative AI and AI-assisted technologies in the writing process

During the preparation of this work, the authors used OpenAI’s ChatGPT to assist with language polishing of parts of the manuscript and to generate original icons for the graphical abstract. The authors carefully reviewed, edited, and verified all AI-generated suggestions and graphical elements and take full responsibility for the final content of the manuscript.

## Methods

### Method details

#### Overview

Poisson processes are well suited for modeling proliferation and differentiation because such cellular events occur at random times, independently of one another, and with rates that remain approximately constant on short time scales. These properties align with the classical assumptions behind Poisson statistics. Consequently, each process occurring within a cell is modeled as a continuous-time Markov chain with Poisson jump intensities with time-dependent parameters. By enumerating all possible processes involved in the considered system, we obtain Eq. (2).

A single cell has a limited set of possible actions (such as activation, self-renewal or differentiation). We assume that the evolution of the entire system is the sum of independent Markov processes corresponding to each type of action. The behavior of an individual cell cannot be directly measured; experimental measurements are performed on groups of cells, tissues, or similar aggregates. This is the level at which we aim to describe the system’s evolution. We therefore assume that many cells follow the same stochastic rules and occupy a sufficiently large region.

If *k* is sufficiently large, it becomes natural to describe the system in terms of concentrations rather than discrete cell numbers. In addition, for large populations, the number of events occurring within a fixed time interval is large, which allows us to approximate the scaled Poisson process by an ODE superposed by its stochastic fluctuation dynamics; see Anderson et al (2015) ^[32]^.

This approach leads to the mathematical results described below. Based on the assumptions above, we can fully characterize the evolution of our system. We show that the evolution of the mean values follows the same ordinary differential equation as in the deterministic model; see Lemma 1. The main advantage of the approach employed in this manuscript is the description of the evolution of the variance in the stochastic model; see Lemma 2. Notably, the variance equation contains the term 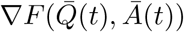, which depends only on the deterministic part of the solution. In this section/appendix we adopted an approach presented by Ethier et al (2009) ^[7]^.

The final result presented here is the transformation of the dynamics from the (*Q, A*)-space to the (*X, Y*)-space. Our analysis operates on two levels. On one hand, the underlying biological processes act on two distinct cell populations, a qNSCs *Q*(*t*) and an aNSCs *A*(*t*). On the other hand, experimental measurements are typically reported in terms of the total number of cells, *X*(*t*) = *Q*(*t*) + *A*(*t*), and the fraction of active cells, 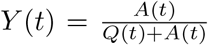. The mathematical approach adopted here allows us to perform a coordinate transformation from (*Q*(*t*), *A*(*t*)) to (*X*(*t*), *Y* (*t*)). This transformation is required both to estimate the parameters of the model and to express the resulting dynamics in the variables corresponding to experimentally observable quantities.

#### Construction of Stochastic NSC Model

Let *Q*^(*k*)^(*t*) and *A*^(*k*)^(*t*) be the number of quiescent and active NSCs at time *t* with initial population

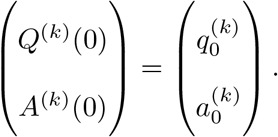

Let 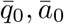 be positive real numbers and let us assume that initial data are described by Gaussian distribution with covariance matrix *U*, i.e. as *k* → ∞, it holds

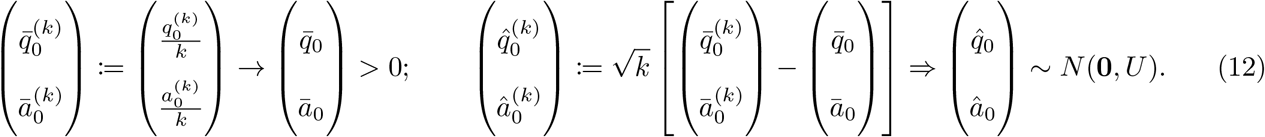

Define

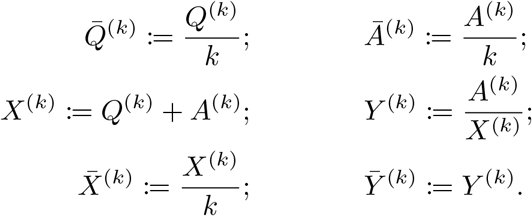

We analyze the system and prove the results on the following class of regulatory mechanism for activation rate *r*(*t*), proliferation rate *p*(*t*), and probability of self-renewal *b*(*t*) such that they are all regulated by 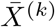. Other regulatory scenarios can be considered in a similar way. Let *r*_0_ *>* 0, *p*_0_ *>* 0, *b*_0_ ∈ (0, 1), the time-dependent parameters are defined by

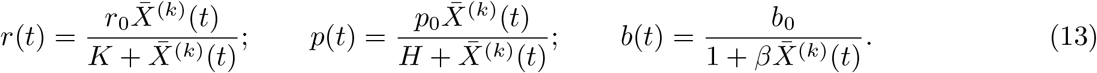

There are three events in this cell proliferation system, activation (ac.), self-renewal (re.), and differentiation (di.). Let *P*_*ac*._, *P*_*re*._, and *P*_*di*._ be three independent unit-rate Poisson process corresponding to three events, we follow the approach by Ethier et al (2009) ^[7]^ and construct the stochastic dynamics as follows.

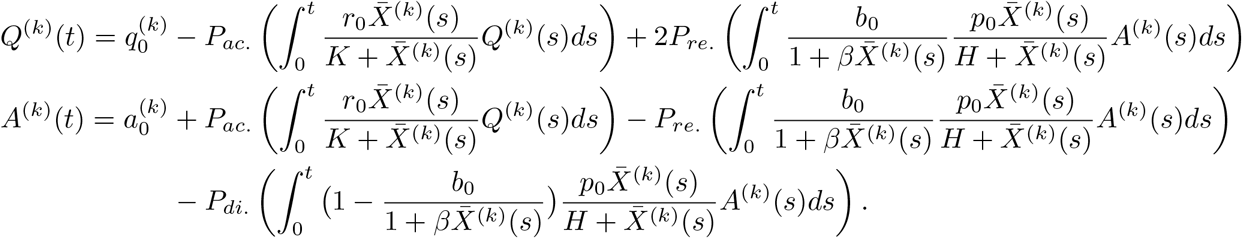

Other scenarios of regulatory mechanisms presented in Eq. (8) can be approached similarly. Furthermore, the cases of parameter combinations considered in Fig. 2C and Fig. 3A can be recovered by setting *K* = 0 or *β* = 0 or *H* = 0.

#### Functional Law of Large Numbers

##### Lemma 1.

*As k* → ∞,

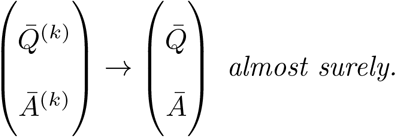

*Where* 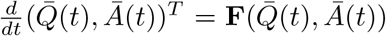 *is an autonomous system of ODEs with initial condition* 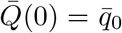 *and* 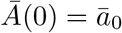. *For the feedbacks in Eq*. (13), **F** *is defined by*

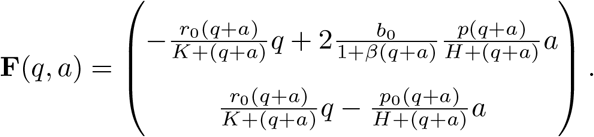

*Proof*. We verify conditions of ^[7]^ Theorem 2.1, Chapter 11 from Ethier et al (2009). The three events in the system described in (2) and Section 4 correspond to the following changes in the population:

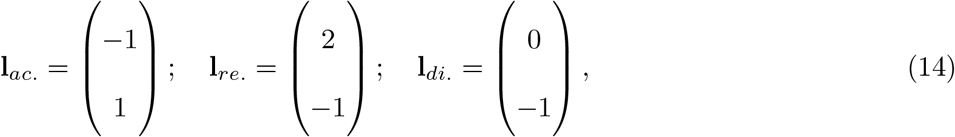

called the stoichiometric vectors. Corresponding rates are equal to

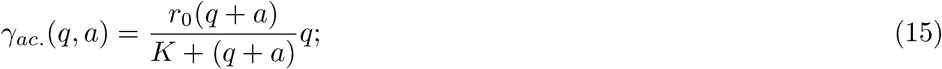

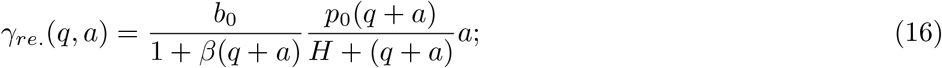

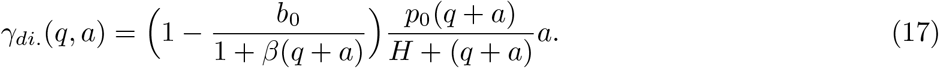

Let *A* = *{ac*., *re*., *di*.*}* be the set of all events in the system. Since for each *α* ∈ *A, γ*_*α*_ is continuous on [0, ∞)^2^, we have for each compact set *K* ⊂ [0, ∞)^2^,

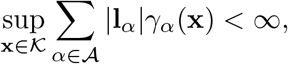

as the sum is taken over a finite set *A*, and where we used | · | to denote the *l*^1^-norm. We now rewrite **F** as

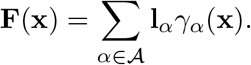

Since each *γ*_*α*_ is continuously differentiable, **F** is locally Lipschitz. Therefore, for each compact *K* ⊂ [0, ∞)^2^, there exists *M*_*K*_ *>* 0 such that for all **x, y** ∈ *K*,

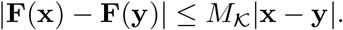

Then the assumptions of ^[7]^ Theorem 2.1, Chapter 11 from Ethier et al (2009) are satisfied. The application of this theorem yields the desired result.

##### Theorem 1.

*Let D be the space of càdlàg functions (Skorokhod space) and define* **g** : *D*^2^ → *D*^2^ *such that*

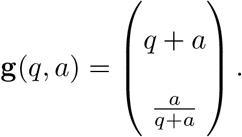

*As k* → ∞,

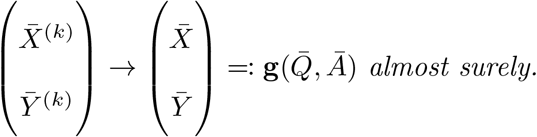

*Proof*. We define **g** : *D × D* → *D × D* such that for all *x, y* ∈ *D*,

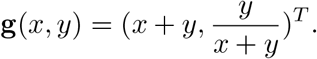

Usually, the addition operator is not continuous on space *D* equipped with Skorokhod metric *d*_∞_. That is, there exists *x*_*n*_ → *x* and *y*_*n*_ → *y* in *D*, but *x*_*n*_ + *y*_*n*_ ↛ *x* + *y*; see^[46]^ Example 3.3.1 from Whitt et al (2002). Nonetheless, if both *x* and *y* are continuous, we do have lim_*n*→∞_(*x*_*n*_ + *y*_*n*_) = *x* + *y*. To see this, suppose *x* and *y* are continuous. We show that for any fixed *T >* 0, *x*_*n*_ → *x* in the sup norm restricted on [0, *T*], denoted ∥ · ∥_*T*_. Since *x*_*n*_ → *x* in *d*_*T*_ (Skorokhod norm restricted on [0, *T*]), there exists a sequence of increasing homeomorphisms *λ*_*n*_ on [0, *T*] such that as *n* → ∞,

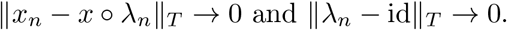

Since continuous function is uniformly continuous on compact sets, there exists a constant *C* such that

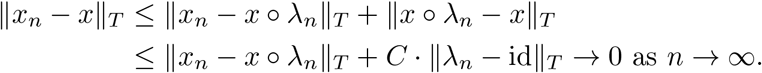

Therefore, both *x*_*n*_ and *y*_*n*_ converge to their limit in ∥ · ∥_*T*_. Using similar reasoning, one can show for all *T >* 0, as long as *x*(*t*) + *y*(*t*) *>* 0 for all *t*, as *n* → ∞,

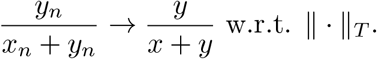

This concludes that **g** is continuous at 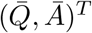 and by the continuous mapping theorem,

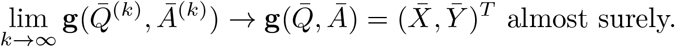

#### Functional Central Limit Theorem

Let Φ be a matrix-valued function satisfying

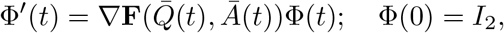

where *I*_2_ ∈ ℝ^2*×*2^ is an identity matrix. Let Ψ = Φ^−1^.

Define a 2*×*3 matrix that encodes the three actions, consisting of the changes (the stoichiometric vectors, see Eq. (14)) in the system and their rates Eq. (15), for the two types of cells 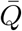 and *Ā*

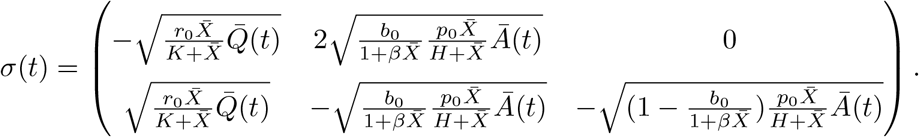

The first column corresponds to activation, the second corresponds to self-renewal, and the final column corresponds to differentiation. Let us define a diffusion matrix Σ(*t*) = *σ*(*t*)*σ*^*T*^ (*t*).

Recall *U* is the covariance function for 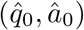 in Section 4. We define the following kernel *K*(*s, t*):

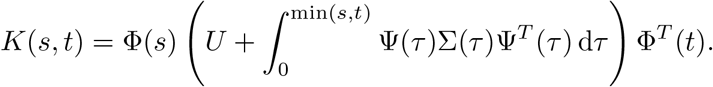

##### Lemma 2.

*As k* → ∞, *we have the following convergence in distribution to a Gauss-Markov process with autocovariance function K*.

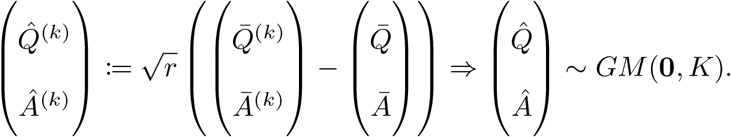

*The variance function satisfies*

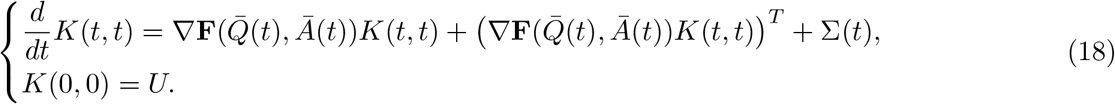

*Proof*. Once more, we prove this lemma by verifying the assumptions of ^[7]^ Theorem 2.3, Chapter 11 by Ethier et al (2009). Since for each *α* ∈ *A, γ*_*α*_ is continuous on [0, ∞)^2^ and number of elements in *A* is finite, we have for each compact set *K* ∈ [0, ∞)^2^,

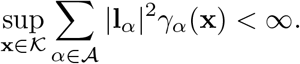

Since *γ*_*α*_’s are continuously differentiable, conditions of ^[7]^ Theorem 2.3, Chapter 11 from Ethier et al (2009) are satisfied. Hence,

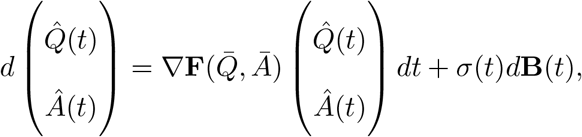

where **B**(·) is a 3−dimensional standard Brownian motion and the diffusion coefficient is

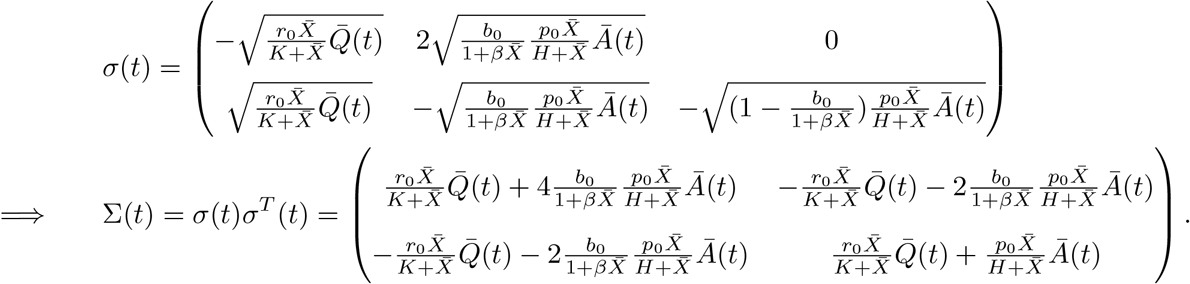

To demonstrate the FCLT dynamics is indeed the claimed Gauss-Markov process, we apply results from^[31]^ Chapter 5.6 of Karatzas et al (2014). The autocovariance function is

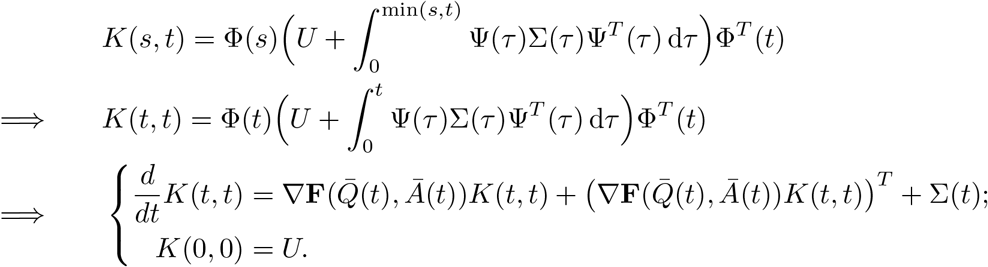

##### Theorem 2.

*Let* **g** : ℝ^2^ → ℝ^2^ *such that*

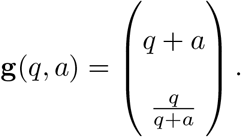

*As k* → ∞, *we have the the following convergence in distribution*

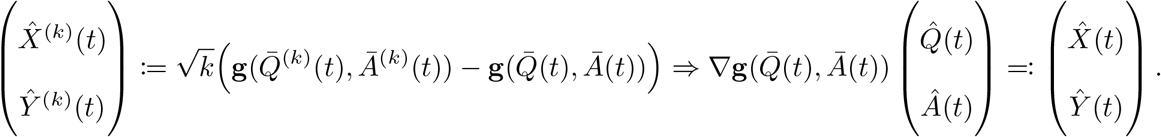

*Proof*. We apply the delta method, see Doob et al (1935) ^[47]^. Fix a time *t* ≥ 0, then we have as *k* → ∞,

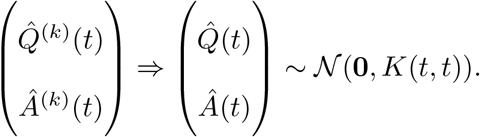

By the definition of 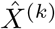 and *Ŷ*^(*k*)^, we apply the delta method^[47]^ to conclude that as *k* → ∞,

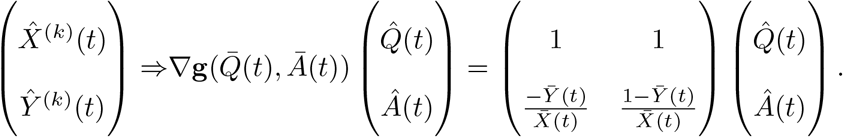

### Quantification and statistical analysis

#### Data preprocessing and parameter estimation

Since our stochastic method relies on a Gaussian assumption, whereas the experimental data exhibit slight skewness, we focus the analysis on the densest region of the data. The full preprocessing and fitting procedure is illustrated in Suppl. Fig. S1A. To this end, we first fit a log-normal distribution (blue curve) to the original data points (shown as large black dots) in order to identify the region of highest data density. The log-normal distribution is then truncated to its main body (red curve) by symmetrically cutting the distribution about its mode. The truncation bounds are defined as the points at which the log-normal probability density function falls below a prescribed threshold. A Gaussian distribution is subsequently fitted to this truncated region (dotted green curve). Filtered data points are shown as smaller red dots; points retained by the filtering therefore appear as red dots with a thick black outline, whereas points excluded by the approximation remain as black dots only.

The resulting Gaussian mean (blue dot) and standard deviation are then used for parameter estimation by minimizing the Bhattacharyya distance between the data-derived and model-predicted Gaussian distributions. For two normal distributions

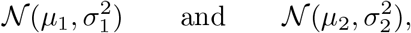

the Bhattacharyya distance is

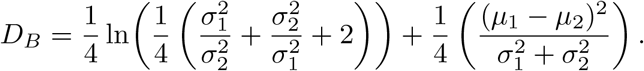

This was used as an objective function, which was minimized using the fmincon routine in MathWorks MATLAB R2022a. The minimization was performed 50-100 times starting from different random initial guesses, and the parameter set with the smallest value of the minimized objective function out of all runs was selected.

#### Model selection

Model selection was carried out by assigning each candidate model a score that balances the quality of the fit against model complexity, thereby reducing the risk of overfitting. A widely used criterion for this purpose is the Akaike Information Criterion (AIC), which assumes a Gaussian error structure based on the Residual Sum of Squares. To account for bias in small sample sizes, Banks et al (2017) ^[48]^ further defined the corrected AIC. In our case, because parameter estimation was performed by minimizing the Bhattacharyya (BHT) distance between model-predicted and empirical distributions, standard AIC scores are not directly applicable. We therefore employed a Bhattacharyya-distance-based analogue of AIC,

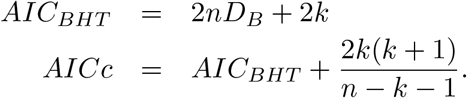

Here, *k* denotes the number of parameters being estimated, *n* is the number of data points, and *D*_*B*_ corresponds to the minimum value of the objective function. Models with lower score values are preferred, as they achieve a better trade-off between fit accuracy and parsimony. The absolute AICc value is not meaningful in isolation; it is the relative differences between models that determine their comparative performance. These differences are expressed as

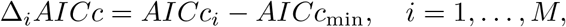

where (AICc_min_) denotes the smallest AICc among the *M* models evaluated. Although only the best model has (Δ_*i*_ AICc = 0), models with small (Δ_*i*_ AICc) values may still be considered competitive. To provide a more quantitative assessment of relative model support, we compute Akaike weights, which estimate the probability that a given model is the most appropriate among the set. The weights are defined as

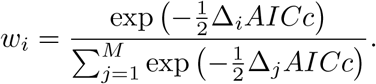

The BHT-adapted AICc and Akaike weights values are shown in Fig. 3B.

