## Supplemental Information for "Harnessing biological variability for mechanistic inference: a stochastic framework applied to neural stem cell dynamics"

### Contents

S1 Validation of prediction intervals using stochastic simulations s1

S2 The existence of a solution to the mean-field ODE s1

S3 Heuristic Derivation for Diffusion Approximation s3

### S1. Validation of prediction intervals using stochastic simulations

Here we present additional figures supporting the main text. Panel A schematically illustrates the data preprocessing and filtering procedure, which is described in detail in the STAR Methods section (Data preprocessing and parameter estimation).
In Panels B and C, rather than displaying the deterministic ODE solution (mean or expected value) together with the prediction intervals as in the main text, we show only the prediction intervals alongside results from stochastic simulations based on 200 independent runs (Fig. S1B and S1C). This representation highlights the close correspondence between the prediction intervals obtained with our method and the variability observed in explicit stochastic trajectories.

### S2. The existence of a solution to the mean-field ODE

**Lemma 1.** *Existence of a global solution to the ODE for the mean value.*

*Proof.* The existence of a local-in-time solution follows from the Picard–Lindelöf theorem. To establish global existence, we define an invariant domain $\mathcal{M}$.
There are two cases to consider:
1. If the initial conditions are equal to zero, i.e. if $(Q_0, A_0) = (0, 0)$, then the solution exists globally in time and it is equal to $(0, 0)$.
2. If at least one of the initial conditions is positive, then the sum of them is also positive, i.e. $X_0 > 0$. In this case, we can use a coordinate transformation from $(Q(t), A(t))$ to $(X(t), Y(t))$, where $X(t) = Q(t)+A(t)$ and $Y(t) = \frac{A(t)}{Q(t)+A(t)}$. Under this transformation, the original system of equations

$$\frac{d}{dt}\begin{pmatrix} Q \\ \\ A \end{pmatrix} = \begin{pmatrix} -\frac{r_0(Q+A)}{K+(Q+A)}Q + 2\frac{b_0}{1+\beta(Q+A)}\frac{p_0(Q+A)}{H+(Q+A)}A \\ \\ \frac{r_0(Q+A)}{K+(Q+A)}Q - \frac{p_0(Q+A)}{H+(Q+A)}A \end{pmatrix} \tag{1}$$

transforms into equivalent system:

$$\frac{d}{dt}\begin{pmatrix} X \\ \\ Y \end{pmatrix} = \begin{pmatrix} \left(2\frac{b_0}{1+\beta X} - 1\right)\frac{p_0 X}{H+X}YX \\ \\ \left(\frac{r_0 X}{K+X}(1-Y) - \frac{p_0 X}{H+X}Y\right) - \left(2\frac{b_0}{1+\beta X} - 1\right)\frac{p_0 X}{H+X}Y^2 \end{pmatrix} \tag{2}$$

Note that the solution $X(t)$ is positive for all $t \in [0, \infty)$. On the other hand $X(t)$ is bounded from above by the maximum of its initial condition $X_0$ and $\tilde{X} = \frac{2b-1}{\beta}$, because $2\frac{b}{1+\beta\tilde{X}} - 1 = 0$. Let us denote this

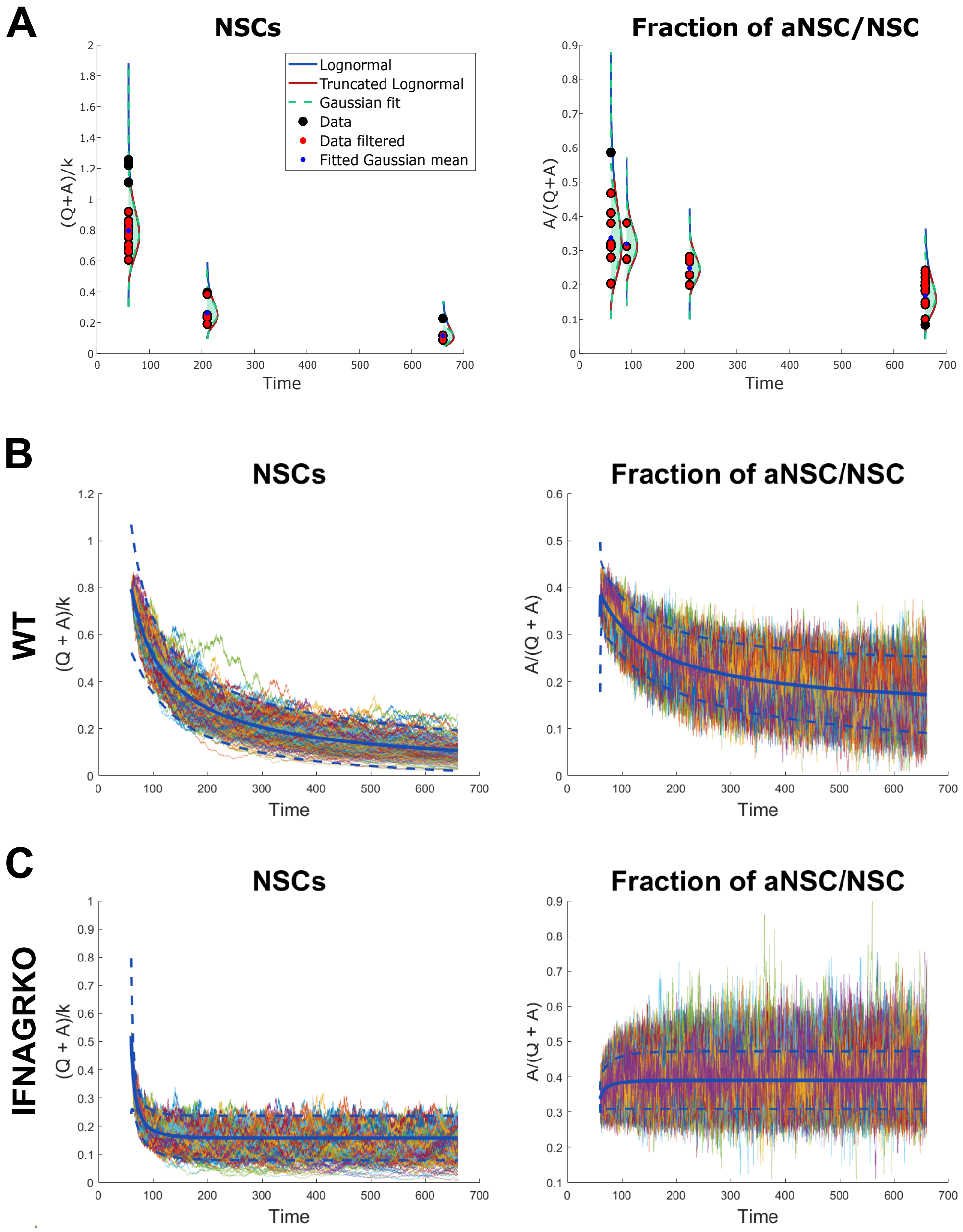


**Figure S1: Stochastic simulations.** In panel **(A)**, we present the data used to calibrate the model. Panels **(B)** and **(C)** present results for wild-type and IFNAGRKO, respectively. The solid blue line represents the mean value and the blue dashed lines indicate the bounds of the prediction interval. All other colored curves represent the results of stochastic simulations.

maximum by $X^*$. Then for all $X \in (0, X^*)$, it holds

$$\begin{aligned}
\frac{d}{dt}Y(t)\Big|_{Y=0} &= \Big(\Big(\frac{r_0 X}{K+X}(1-Y) - \frac{p_0 X}{H+X}Y\Big) - \big(2\frac{b_0}{1+\beta X} - 1\big)\frac{p_0 X}{H+X}Y^2\Big)\Big|_{Y=0} = \frac{r_0 X}{K+X} > 0 \\
\frac{d}{dt}Y(t)\Big|_{Y=1} &= \Big(\Big(\frac{r_0 X}{K+X}(1-Y) - \frac{p_0 X}{H+X}Y\Big) - \big(2\frac{b_0}{1+\beta X} - 1\big)\frac{p_0 X}{H+X}Y^2\Big)\Big|_{Y=1} \\
&= -\frac{p_0 X}{H+X} - \big(2\frac{b_0}{1+\beta X} - 1\big)\frac{p_0 X}{H+X} = -\frac{p_0 X}{H+X}\big(1 + 2\frac{b_0}{1+\beta X} - 1\big) \\
&= -2\frac{p_0 X}{H+X}\frac{b_0}{1+\beta X} < 0
\end{aligned}$$

This implies that the solution $(X(t), Y(t))$ starting from $\mathcal{M} = [0, X^*] \times [0, 1]$ does not leave this domain.

□

### S3. Heuristic Derivation for Diffusion Approximation

In this section, we provide heuristic derivation following Anderson et al (2015) [?]. Denote $\bar{\mathbf{N}}^{(k)} = (\bar{Q}^{(k)}, \bar{A}^{(k)})^T$, $\bar{\mathbf{N}} = (\bar{Q}, \bar{A})^T$, and $\hat{\mathbf{N}}^{(k)} = (\hat{Q}^{(k)}, \hat{A}^{(k)})^T$. Denote $\hat{\mathbf{N}}(0) = \hat{\mathbf{n}}$, then

$$\begin{aligned}
\bar{\mathbf{N}}^{(k)}(t) &= \bar{\mathbf{N}}^{(k)}(0) + \int_0^t \mathbf{F}(\bar{\mathbf{N}}^{(k)}(\tau))\,\mathrm{d}\tau \\
&\quad + \sum_{\alpha\in\mathcal{A}} \frac{1}{k}\mathbf{l}_\alpha \left(P_\alpha\left(k\int_0^t \gamma_\alpha(\bar{\mathbf{N}}^{(k)}(\tau))\,\mathrm{d}\tau\right) - \int_0^t k\gamma_\alpha(\bar{\mathbf{N}}^{(k)}(\tau))d\tau\right) \\
\bar{\mathbf{N}}(t) &= \bar{\mathbf{N}}(0) + \int_0^t \mathbf{F}(\bar{\mathbf{N}}(\tau))\,\mathrm{d}\tau.
\end{aligned}$$

Therefore,

$$\begin{aligned}
\hat{\mathbf{N}}^{(k)}(t) &= \hat{\mathbf{N}}^{(k)}(0) + \int_0^t \sqrt{k}\left(\mathbf{F}(\bar{\mathbf{N}}^{(k)}(\tau)) - \mathbf{F}(\bar{\mathbf{N}}(\tau))\right)\,\mathrm{d}\tau \\
&\quad + \sum_{\alpha\in\mathcal{A}} \frac{1}{\sqrt{k}}\mathbf{l}_\alpha \left(P_\alpha\left(k\int_0^t \gamma_\alpha(\bar{\mathbf{N}}^{(k)}(\tau))\,\mathrm{d}\tau\right) - \int_0^t k\gamma_\alpha(\bar{\mathbf{N}}^{(k)}(\tau))d\tau\right).
\end{aligned}$$

For large $k$, we have

$$\begin{aligned}
&\hat{\mathbf{N}}^{(k)}(0) \approx \hat{\mathbf{n}}, \\
&\sqrt{k}\left(\mathbf{F}(\bar{\mathbf{N}}^{(k)}(\tau)) - \mathbf{F}(\bar{\mathbf{N}}(\tau))\right) \approx \nabla\mathbf{F}(\bar{\mathbf{N}}(\tau))\hat{\mathbf{N}}(\tau), \\
&\frac{1}{\sqrt{k}}\left(P_\alpha\left(k\int_0^t \gamma_\alpha(\bar{\mathbf{N}}^{(k)}(\tau))\,\mathrm{d}\tau\right) - \int_0^t k\gamma_\alpha(\bar{\mathbf{N}}^{(k)}(\tau))d\tau\right) \approx B_\alpha\left(\int_0^t \gamma_\alpha(\bar{\mathbf{N}}^{(k)}(\tau))\,\mathrm{d}\tau\right),
\end{aligned}$$

where $B_\alpha$'s are independent standard Brownian motions associated with actions $\alpha \in \mathcal{A}$. The last approximation is a consequence of the Poisson FCLT [? ? ?]. Hence,

$$\hat{\mathbf{N}}^{(k)}(t) \approx \hat{\mathbf{n}} + \int_0^t \nabla\mathbf{F}(\bar{\mathbf{N}}(\tau))\hat{\mathbf{N}}(\tau)d\tau + \sum_{\alpha\in\mathcal{A}} \mathbf{l}_\alpha B_\alpha\left(\int_0^t \gamma_\alpha(\bar{\mathbf{N}}^{(k)}(\tau))\,\mathrm{d}\tau\right).$$
